## Supplementary material for "Modelling Amoebic Brain Infection Caused by *Balamuthia mandrillaris* Using a Human Cerebral Organoid": Legend of supplemental figures

**Supplementary data**

**Figure S1.** **Cytopathologic effect of *B. mandrillaris* trophozoites on human cerebral organoids.** A 16-day coculture of *B. mandrillaris* trophozoites with human cerebral organoids was subjected to H&E staining. (A) Trophozoites of *B. mandrillaris* were observed inside the cerebral organoid, in which the cells formed clumps (white insets). (B and C) Higher magnification of insets in panel A shows trophozoites in the cell clump in the upper and lower inset of panel A, respectively. (D and E) Zoomed-in images of the insets in panels B and C, respectively. Yellow arrows indicate trophozoites with a round shape with nuclei in some cells. F. Noninfected cerebral organoids showed well-defined nuclei without cell clumps. Scale bars in panels A-C = 100 µm; scale bars in panels D-E = 50 µm.

**Figure S2.** **Coculture of** ***B. mandrillaris* trophozoites with human cerebral organoids. (A)** Representative fluorescence images of cerebral organoids cocultured with trophozoites (scale bars, 200 µm). The cerebral organoids and the trophozoites were differentially labelled using the protein-binding CMFDA (green) and lipid-binding DiD (magenta) fluorescent markers, respectively. **(B)** Microscopy images of cerebral organoids cultured with trophozoites. On the surface, the trophozoites had multiple cytoplasm protrusions with active movement (arrowheads, left panel). Two days later, there were many trophozoites located at a small focal break in the cerebral organoids (arrowheads, right panel). Some trophozoites remained elongated in shape (asterisks, right panel), while most were oval (arrowheads, right panel). Scale bars, 100 µm.
