## Supplementary figures and images for "Modelling Amoebic Brain Infection Caused by *Balamuthia mandrillaris* Using a Human Cerebral Organoid"

### Supplemental figure 1

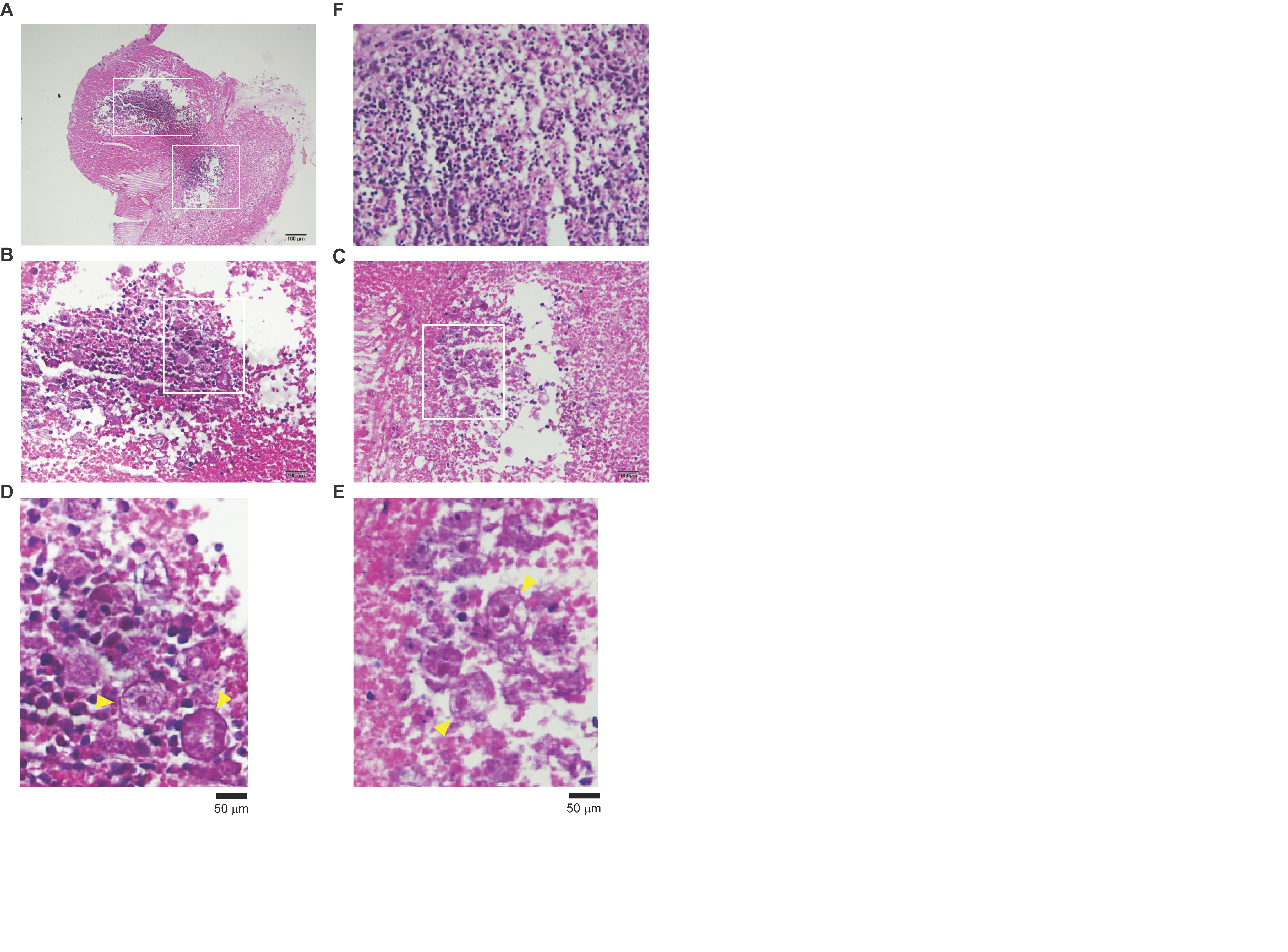

### Supplemental figure 2

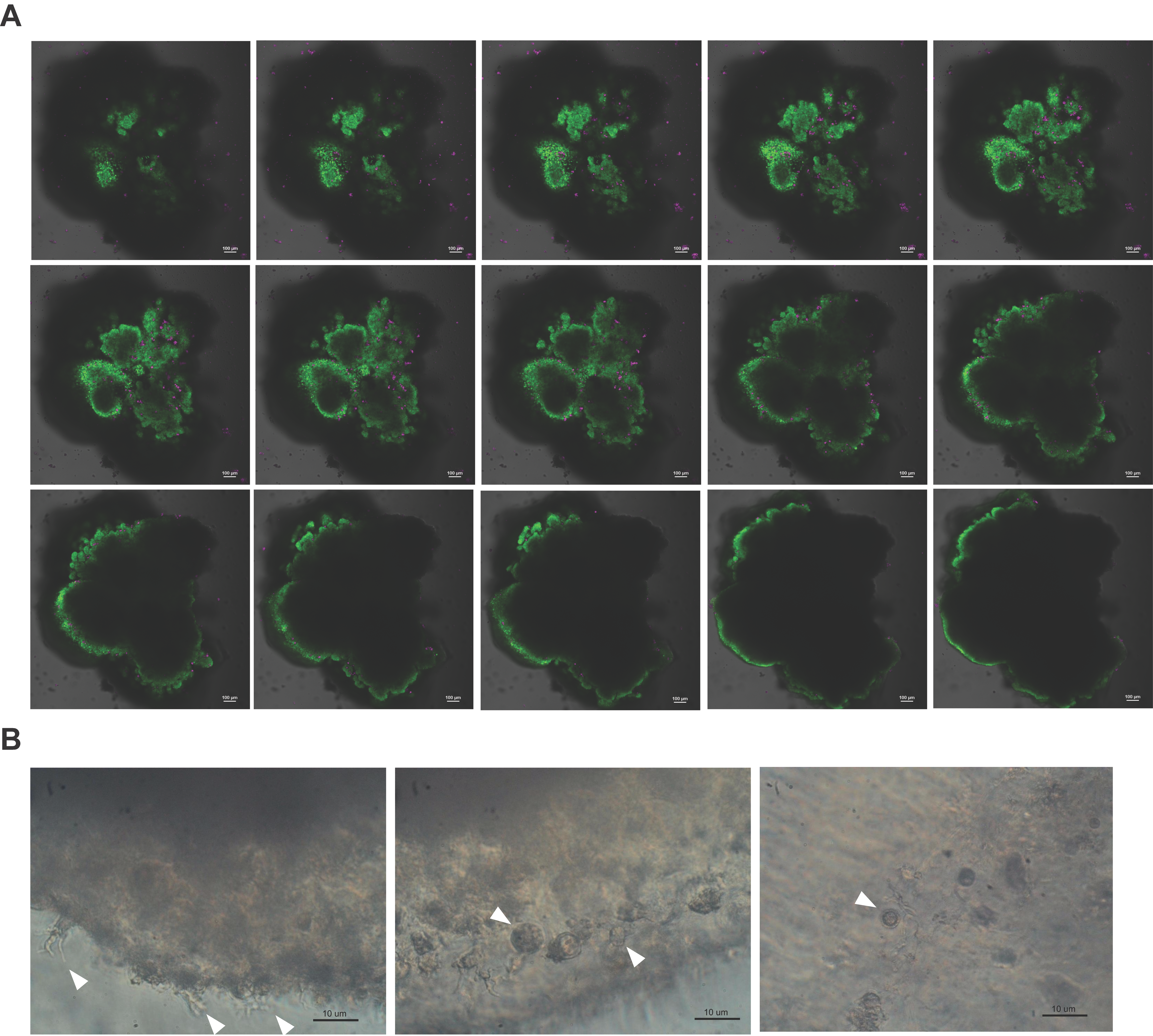
